# Nurture over nature in new *Micromonospora* spp. strains’ metabolism: multi-omics analyses reveal fermentation conditions as the dominant driver over phylogeny

**DOI:** 10.1101/2025.11.08.687343

**Authors:** Jia-Rui Han, Shuai Li, Wen-Hui Lian, Lu Xu, Li Duan, Jia-Ling Li, Guo-Yuan Shi, Qi-Chuang Wei, Mukhtiar Ali, Wen-Jun Li, Lei Dong

**Author notes:** Corresponding authors: Mukhtiar Ali, Wen-Jun Li; Lei Dong.

## Abstract

The genus *Micromonospora*, a key member of the actinomycetes, has demonstrated considerable potential for natural product biosynthesis. In this study, we isolated 15 *Micromonospora* spp. strains from desert soil and marine sediment samples, eight of which represent four novel species. To explore the biosynthetic capacity of this genus, we performed an integrated analysis of *Micromonospora* reference genomes. Pan-genomic analysis further unveiled the core biosynthetic characteristics of the genus responsible for producing terpenes and polyketides. Further multi-omics investigation, combining genomic and metabolomic data, uncovered a positive correlation between phylogenetic relationships and biosynthetic potential, alongside a decoupling of metabolic profiles. Notably, metabolomic findings emphasized the dominant influence of culture conditions on the expression of biosynthetic capabilities. Overall, our study provides a comprehensive elucidation of the biosynthetic potential of the genus *Micromonospora* and highlights the value of investigating novel strains and applying diverse cultivation strategies in natural product discovery.

**Importance:** Our study provides a comprehensive genomic and metabolomic elucidation of the significant biosynthetic potential within the genus *Micromonospora*. It reveals a core biosynthetic capacity for terpenes and polyketides that is phylogenetically linked, whereas the resulting natural product repertoire is subject to strong modulation by cultivation conditions. These findings underscore the critical importance of exploring novel species and employing diverse cultivation strategies to unlock the full potential of microbial resources for natural product discovery.

## Introduction

Throughout history, microbial natural products have been extensively utilized as precursors for antibiotics, antitumor compounds, and various other pharmaceuticals (1, 2). Certainly, these natural products continue to play a significant role in modern medicine and remain a primary source for the development of novel small molecule drugs to address global antibiotic resistance and other public health challenges (3). Approximately 70% of the natural drugs currently in use are derived from *Actinobacteria* (4). As a member of Actinobacteria, the genus *Micromonospora* has been widely reported to produce natural products with unique chemical diversity and enormous therapeutic potential, representing an untapped resource for drug and drug-lead discovery (5). The genus Micromonospora was originally proposed by Ørskov in 1923 (6). Members of this genus are Gram-stain positive, spore-forming aerobic *Actinobacteria* that possess unique morphological characteristics such as a single spore attached to short substrate hyphae (7). Members of *Micromonospora* are widely distributed in nature and has been isolated and cultivated from various samples, including soil (8–10), water (11–13), plants (14, 15) and animals (16, 17), and the genus *Micromonospora* is currently the second-largest group within the phylum *Actinomycetota* (https://lpsn.dsmz.de/genus/micromonospora). Their metabolites exhibited excellent antibacterial and anticancer activities, particularly compounds belonging to the macrolide (18, 19), peptide (20, 21), aminoglycoside (8, 22), and ansamycin classes (23, 24). Therefore, it is essential to systematically investigate the biosynthetic potential of this genus under different ecosystems.

However, according to reports, the past two decades have witnessed a significant decline in the discovery of bioactive secondary metabolites with novel structures derived from *Micromonospora* (5, 25). Evidently, the golden age of antibiotic discovery (1950–1960) has ended (26), accompanied by a steady decrease in the proportion of new species and compounds obtained from common environmental samples. Moreover, traditional natural product discovery has relied heavily on culturing microorganisms in the laboratory. However, it is estimated that only a small fraction (1-10%) of environmental microbes can be cultured using standard techniques and the majority of microbial diversity remains unknown(27). While the discovery of new secondary metabolites from *Micromonospora* has decreased, exploring new habitats could lead to the identification of novel microbial species, which may in turn uncover new chemical diversity and secondary metabolites (28, 29). Therefore, the recent focus of researchers is increasingly shifting toward extreme habitats such as deserts, deep seas, volcanoes, and polar regions, to discover new microbial resources that can produce structurally novel and promising natural products (29–31).

In recent years, the development and widespread use of advanced biological detection and analysis tools have enabled researchers to discover that even environments previously considered “forbidden zones of life”, contain a large, untapped microbial diversity reservoir (32). Moreover, undertaking the simultaneous tasks of extracting unspecified bacterial strains from natural environmental samples, screening the metabolite production of those strains, isolating novel chemical compounds, and elucidating the underlying biosynthetic pathways is undoubtedly similar to “needle in a haystack” (33). Employing advanced non-targeted screening analysis and multi-omics techniques to investigate the biosynthetic potential, combined with targeted screening of microbial strains and advanced processing methods, can substantially reduce the burden of exploring novel strains with the biosynthetic potential (34).

This study focuses on the entire genus *Micromonospora*. Through comprehensive genome and pangenome analyses, we investigated the biosynthetic potential of this genus at the genomic level and characterized the distribution patterns of biosynthetic gene clusters. Furthermore, multi-omics analyses were conducted on *Micromonospora* strains isolated from marine and desert samples in this research to explore the relationships among phylogeny, biosynthetic potential, and metabolic profiles. Through this approach, we aim to efficiently uncover bioactive natural products from *Micromonospora* isolates and establish a multi-omics analytical framework for systematically exploring the biosynthetic potential of wild-type strains.

## Results

### Phylogenetic relationship

The 15 isolates were classified into the genus *Micromonospora* based on the 16S rRNA sequence similarity comparisons and phylogenetic analyses (**Table S1**, **Fig. S1**). The results showed that the 15 isolates were clustered closely with the validly published strains of the genus *Micromonospora*. Moreover, whole genome sequencing was conducted for the isolates and the average nucleotide identity (ANI) was calculated between the genomes of 15 isolates and *Micromonospora* type strains and the highest ANI values of the 15 isolates and their corresponding type strains were recorded as shown in **Table S2**. In addition, it was found that the ANI values of 8 species, including SYSU B006, SYSU D00602, SYSU D01140, SYSU D00755, SYSU D00756, SYSU D01135, SYSU D00963, and SYSU D00964, were all below 95%, which is the delineation threshold value for a novel species definition (35, 36). Moreover, digital DNA-DNA hybridization (dDDH) was performed to measure the degree of genetic similarity between the pools of DNA sequences among the eight isolates (with ANI <95%) as shown in **Table S3**. The results showed that all the dDDH values were below 70% which is also a threshold defined for a novel species (35, 36). As described by Han et al. (37), through polyphasic taxonomic characterization of the eight potential novel strains, four novel species were formally proposed: SYSU B006 represents *Micromonospora marincola* sp. nov.; SYSU D00602 and SYSU D01140 represent *Micromonospora desertarenae* sp. nov.; SYSU D00755, SYSU D00756 and SYSU D01135 represent *Micromonospora aridisoli* sp. nov.; and SYSU D00963 and SYSU D00964 represent *Micromonospora xinjiangensis* sp. nov.. The gene sequences of all isolated strains have been deposited in the NCBI database. See **Table S5** for deposition details.

### Elucidating biosynthetic resources in the genus *Micromonospora*

Phylogenetic relationships and an overview of biosynthetic resources in the genus *Micromonospora* were shown in **Fig. 1A**. Moreover, the whole genomes of *Micromonospora* type strains obtained in this study were analyzed with the genomes of the novel *Micromonospora* strains isolated and cultured to explore further the desired niche associated with the novel strains. All the analyzed strains in this study had complete information regarding their isolation sources (**Table S4**). Therefore, based on this information, the strains were categorized into five groups: soil, desert, water, host, and others. The whole genome analysis revealed that the highest strains were recorded in soil group (39 genomes) followed by the host (32 genomes), water (16 genomes), and desert (7 genomes) group, respectively, whereas the lowest strains (2 genomes) were recorded in others group in this study. These results stated that the main sources of *Micromonospora* isolates are soil and host organisms.

**Fig. 1.**
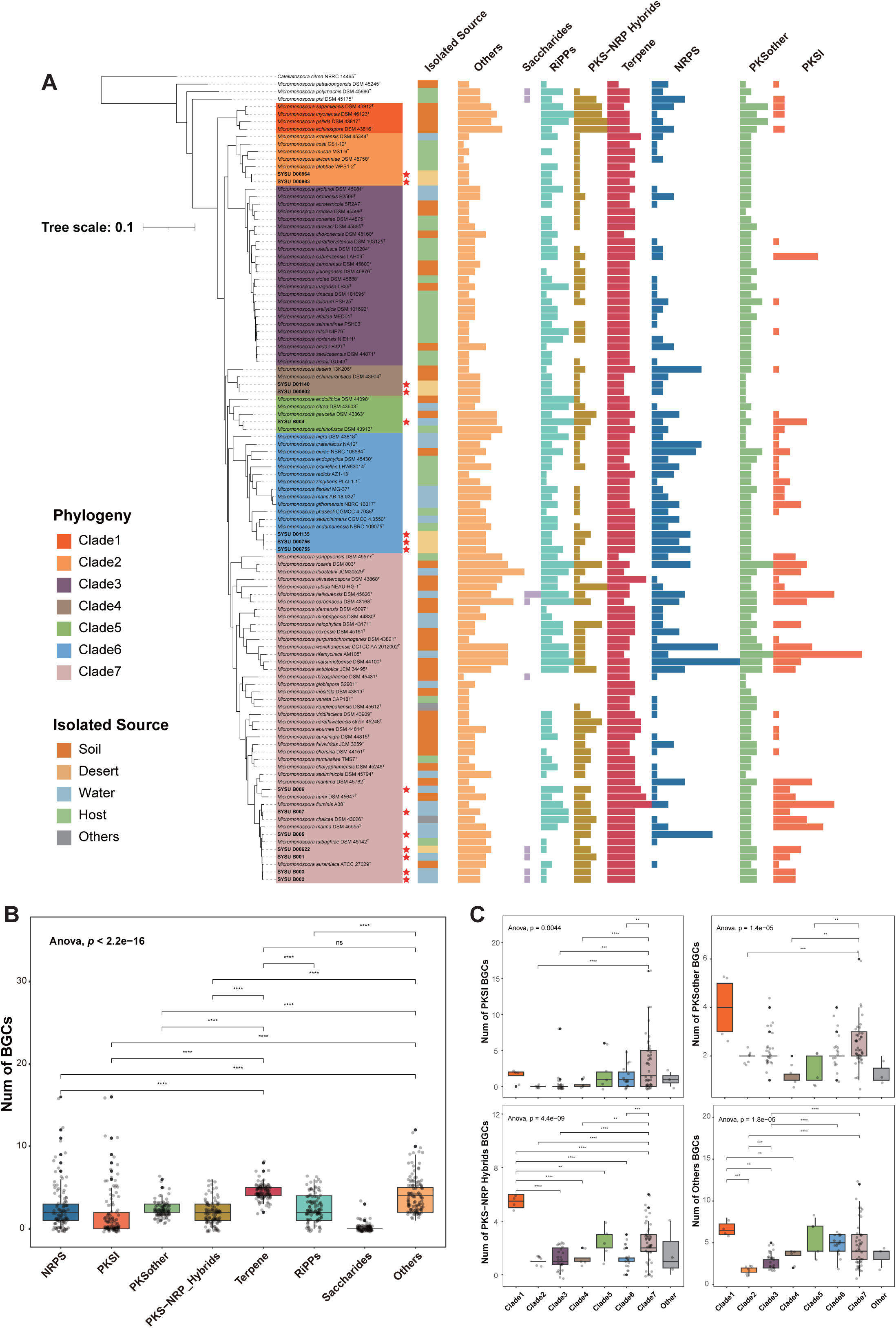
Phylogenetic relationship and overview of biosynthetic potential in the genus *Micromonospora*. **A** The maximum likelihood (ML) phylogram of 107 *Micromonospora* genomes was constructed using the sequences of 120 ubiquitous single-copy proteins, and *Catellatospora citrea* NBRC 14495^T^ (GCA_016862615.1) was used as outgroup. The phylogenetic tree divided the *Micromonospora* strains into 7 clades, and the red star symbols at the outer edges of the isolate names represent strains isolated in this study. The different colors in the right bar chart indicate the source of isolation of the corresponding strain, and the most right combined bar chart reflects the type and amount of BGCs of the corresponding strain. **B** Scatter plot and box plot of BGC counts in genus *Micromonospora* **C** Scatter plot and box plot of PKSI, PKSother, PKS-NRP_Hybrids, Others BGC counts in genus *Micromonospora* based on phylogenetic relationships grouping. Significance levels are as follows: *p* > 0.05 = n.s., *p* < 0.5 = *, *p* < 0.01 = **, *p* <0.001 = ***, *p* < 0.0001 = ****.

The strains were further divided into seven clades based on phylogenetic gene tree topology, annotated numbers, and class of biosynthetic gene clusters (BGCs) identified by antiSMASH and BiG-SCAPE as shown in **Figs. 1A**, **1B**. Biosynthetic diversity evaluation at the clade level was done to investigate the identification of a *Micromonospora* phylogenetic lineage capable of possessing the highest biosynthetic potential with the plethora of BGCs. Therefore, biosynthetic potential annotation for a total of 107 *Micromonospora* genomes was further performed representing the genus *Micromonospora* and the results revealed the identification of 2,106 BGCs. Among all the BGCs, the highest number of Terpene (22.6%) were recorded, followed by RiPPs (12.9%), NRPS (12.7%), PKSother (11.7%), PKS-NRP_Hybrids (9.5%), and PKSI (8.9%). Numerous hybrid-classified BGCs were grouped under “Others” in BiG-SCAPE annotation and constituted a significant proportion (21.2%) as shown in **Fig. 1B**. Further analysis revealed significant differences in the number of BGCs present across strains belonging to different phylogenetic clades (**Fig. 1C**, **S2**). Notably, such differences were more pronounced in BGC categories such as PKSI, PKSother, PKS-NRP_Hybrids, and Others. When the genomes were grouped by isolation sources, no significant differences in the number of BGCs were observed among the groups (**Fig. S3**). These findings suggested a potential correlation between the biosynthetic potential of different strains and their phylogenetic relationships.

Gene Cluster Families (GCFs) composed of BGCs predicted and annotated from 107 *Micromonospora* genomes were analyzed through BiG-SCAPE (**Fig 2A**). The results indicated a strong clustering relationship among Terpene BGCs compared to the other BGCs. Moreover, the results obtained from antiSMASH were organized and it was found that the BGCs annotation in the genus *Micromonospora* showed a low homology (with an average similarity of only 25%) to the encoded known BGCs for natural products based on the available database (**Fig. 2B**). Annotation results revealed that among the total BGCs observed, 442 BGCs (20.1% of the total) were having 0% of the similarity. Moreover, the number of BGCs with a similarity score less than or equal to 50% was 1761 (83.6% of the total). On the other hand, only 52 BGCs (2.5% of the total) exhibited a 100% similarity. These results indicated that the genus *Micromonospora* possesses highly unique biosynthetic resources that are yet to be explored.

**Fig. 2.**
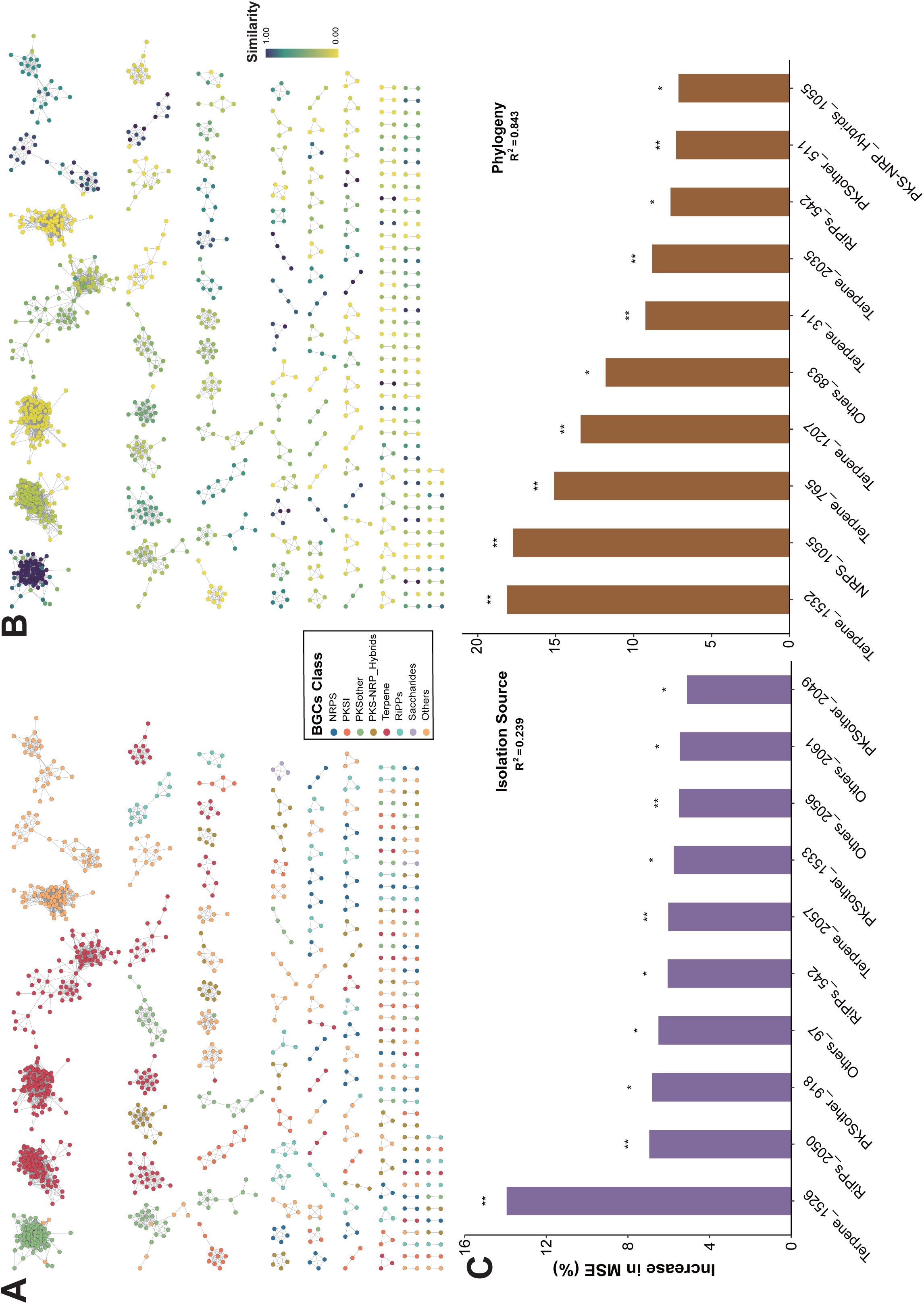
Overview of GCFs in the genus *Micromonospora*. **A** The GCFs composed of BGCs predicted and annotated from 107 *Micromonospora* genomes. Each node in the network represents a BGC, and different colors indicate different classes of BGCs. **B** The GCFs annotated based on similarity to BGCs encoding known natural products in the database. The colors represent the decreasing similarity and increasing novelty of the BGCs. **C** Random Forest (RF) models (5000 trees) constructed based on the distribution of GCFs in the *Micromonospora* isolates, using isolation source and phylogeny as grouping criteria. The bar chart displays the top 10 GCFs with the highest mean predictor importance (percentage of increase of mean square error) in each model. Percentage increases in the MSE (mean squared error) of variables were used to estimate the importance of these predictors, and higher MSE% values imply more important predictors. Significance levels are as follows: *p* < 0.5 = *, *p* < 0.01 = **. MSE, mean squared error.

Random forest models were constructed based on the distribution of *Micromonospora* strains in the GCFs through the grouping criteria of isolation source and phylogeny (**Fig. 2C**). This model was used as an estimator to fit the phylogenetic tree of isolation source and used averaging GCFs to expand the predictive accuracy. The results revealed that among the different classes of biosynthetic potentials, GCFs belonging to the class Terpene exhibited the highest contributions. In addition, the phylogeny group (R^2^=0.843) exhibited higher explanatory power and better fit compared to the isolation source group(R^2^=0.239), indicating that the diversity of the *Micromonospora* phylogeny had a high impact on the diversity of its biosynthetic potential compared to the isolation source.

The 15 strains isolated in this study exhibited significant phylogenetic diversity and occupied five out of the seven clades of the genus *Micromonospora* (**Fig. 1A**). Especially, eight strains from desert sources were distributed across four clades, while the remaining seven strains from marine sources were distributed across two clades. Moreover, it was revealed that the desert-derived isolates displayed greater phylogenetic diversity compared to the marine-derived isolates. In addition, the biosynthetic characteristics of 15 isolates were highly similar to the other members of the genus *Micromonospora*, resulting in the identification of a total of 306 BGCs. The major BGCs categories, including Terpene (22.9%), NRPS (12.1%), PKS-NRP_Hybrids (11.4%), RiPPs (11.1%), PKSother (10.5%), and PKSI (9.2%).

GCFs were further calculated through the incorporation of BGCs from the MIBiG dataset(38). Clustering of the annotated BGCs from 15 isolates resulted in the identification of 3 GCFs containing reference BGCs (**Fig. S4**). The local BGCs form similarity networks with reference BGCs of Loxoribicin A1 and A2 (compounds **13** and **14**), Rakicidins A and B (compounds **17** and **18**), as well as Diazaquinomycins H and J (compounds **15** and **16**). These results indicated that the isolates have the potential to produce natural products with bioactivity and structural similarity to these known compounds.

In summary, the genus *Micromonospora* demonstrated rich phylogenetic diversity and exhibited a remarkable diversity and novelty of biosynthetic resources. The isolated strains in this study, particularly those derived from desert environments, further highlight these characteristics.

### Pan-genome and biosynthetic gene distribution in genus *Micromonospora*

Pan-genome provides insights into genomic variation and differential gene expression profiles related to biosynthesis. Therefore, a pan-genomic analysis of 107 *Micromonospora* genomes was constructed to obtain additional information about the genetic diversity through a pan-genome dataset for the genus *Micromonospora* (**Fig. 3A**). The pan-genomic analysis revealed that the average number of core genes in *Micromonospora* pan-genome dataset was 1,632, while the number of accessory genes was 4,357, and the number of singleton genes was 327. These results showed that the number of accessory genes in pan genome were generally the most abundant in each genome and accounted for an average proportion of 67.0%, while singleton genes were the least abundant with an average proportion of 5.2%. Furthermore, pan-genomic analysis for *Micromonospora* strains grouped by different isolation sources were performed, including soil, desert, water, and host as shown in (**Fig 3B**). As discussed in the previous section, the source analysis revealed that the soil and host groups contained more genomes with 39 and 32 genomes each, respectively. The pan-genome analysis of the two groups revealed that both groups exhibited relatively fewer core genes (with an average value of 1,941 (soil) and 1,937 (host), respectively) and more accessory genes (with average values of 3,828 and 3,855, respectively). On the contrary basis, as discussed in the previous section, water and desert groups had relatively fewer genomes with 16 and 8 genomes each, respectively. The pan-genome analysis for these two groups revealed that water and desert groups had relatively more core genes (with average values of 2,301 and 3,109, respectively) and fewer accessory genes (with average values of 3,277 and 2,687, respectively). Though there were significant differences in the number of accessory and core genes among these four groups (soil, host, water, desert), the number of singleton genes showed no significant difference. These results suggested that the *Micromonospora* genus pan-genome tends to stabilize as more genomes are included with a gradual reduction in core genes (genome present in all the strains) and an increase in accessory genes (genome not present in all strains), while singleton genes remain relatively stable.

**Fig. 3.**
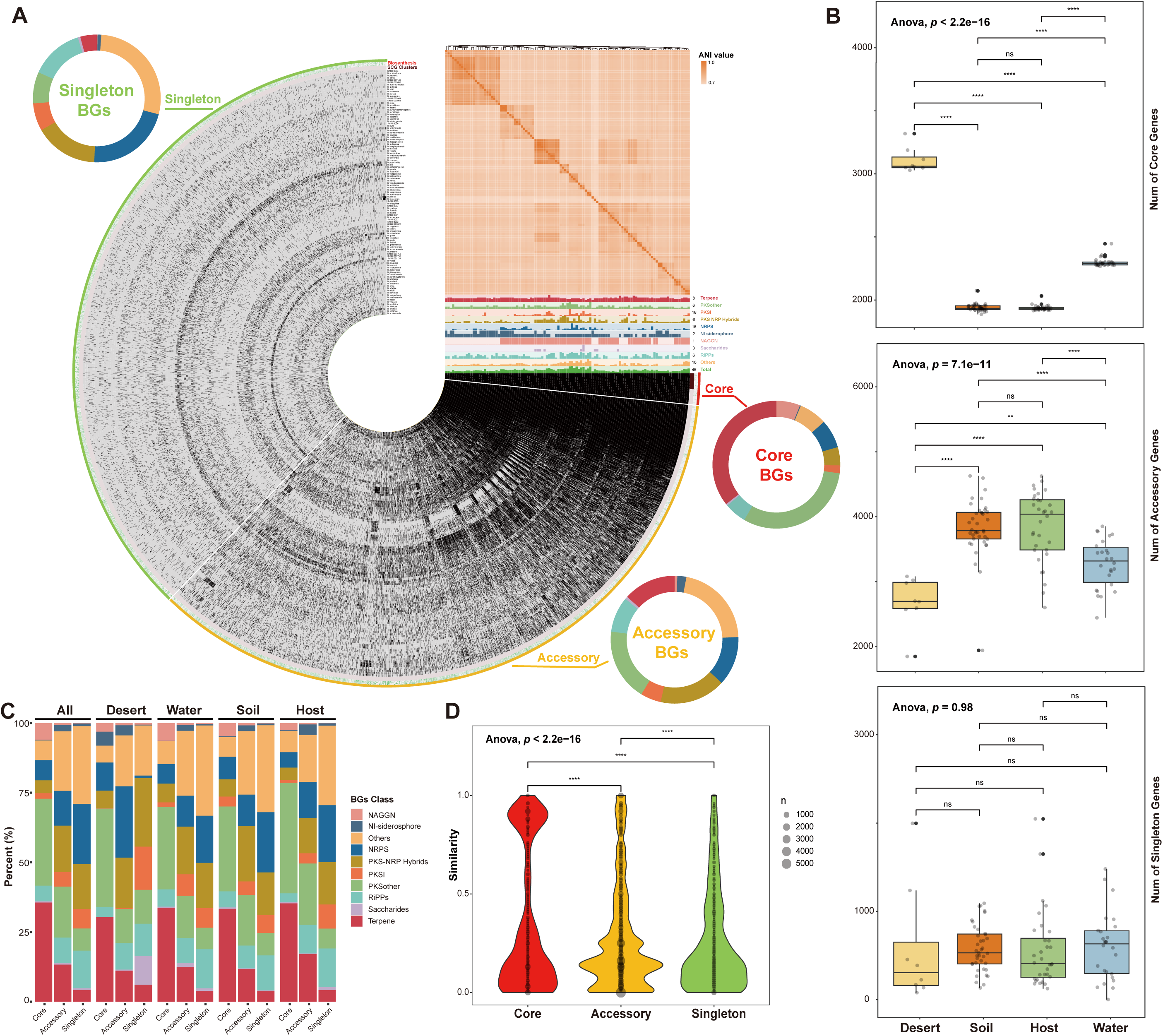
Pan-genome characteristics and distribution of biosynthetic genes in the genus *Micromonospora*. **A** In the circular interface, each layer (gray) represents all genes (black) within an individual genome, along with the distribution of SCG (single-copy core genes) clusters (brown) and BGs distribution (green). For the Bin name, the core region (red) comprises genes present in all 107 *Micromonospora* genomes, the accessory region (yellow) contains genes shared among some genomes, and the singleton region (green) encompasses species-specific genes present in only one genome. The distribution of BGCs in strains is depicted by the histogram in the top right of the circular interface, with the maximum number and classification of each BGC displayed to the right of the histogram. The heatmap above the BGCs histogram shows the ANI values between genomes. The outer BGC doughnut chart provides an overview of the proportion of BG categories within each Bin. **B** Scatter and box plots of the counts of core genes, accessory genes, and singleton genes in the pan-genome datasets constructed for the *Micromonospora* strains based on different isolation sources. **C** Stack plot showing the proportion of the classes of BGs mapped to in the pan-genome datasets and their distribution among different isolation source groups. **D** Violin and bubble plots displaying the similarity of core genes, accessory genes, and singleton genes containing BGs in the pan-genome datasets to those encoding known natural products in the database. Significance levels are as follows: *p* > 0.05 = n.s., *p* < 0.5 = *, *p* < 0.01 = **, *p* <0.001 = ***, *p* < 0.0001 = ****.

The BGC sequences annotated by antiSMASH were mapped to *Micromonospora* pan-genome dataset to further investigate the biosynthetic genes (BGs) distribution (**Fig. 3A, 3C**), which revealed that the BGs classes were mostly aligned with the BGCs annotation results from BiG-SCAPE. Moreover, we separated the two classes, NI-siderophore and NAGGN (N-Acetylglutaminylglutamine amide) from the other classes due to their significant amount and essentiality for strain survival. The analysis revealed that Terpene and PKSother constituted significant proportions of the pan-genome dataset, comprising 35.6% and 31.2%, respectively. The pan-genome analysis revealed that NAGGN BGs showed a higher proportion in core genes (6.0%) compared to accessory genes (0.7%) and singleton genes (0.2%). On the other hand, the NI-siderophore BGs had a relatively higher proportion in accessory genes at 2.3% compared to 0.2% in core genes and 0.9% in singleton genes. As the desert represents an extreme habitat source for all microorganisms, including *Micromonospora* strains, the core genes distribution in the pan-genome dataset from desert soil shows higher proportions of PKSother (35.4% compared to the overall 31.2%), NI-siderophore (5.0% compared to the overall 0.2%), NRPS (10.1% compared to the overall 7.2%), and PKS-NRP_Hybrids BGs (6.3% compared to the overall 4.6%). The novelty of BGs within the pan-genome dataset was shown in **Fig. 3D**, **S5**. Significant differences among the pan-genome datasets were recorded and the results revealed that BGs in the core genes exhibited a relatively higher average similarity of 36.1%, while BGs in the accessory genes had an average similarity of 27.7%. The lowest BGs were recorded in the singleton gene with an average similarity of 25.9%. Despite being conventional sequences shared among *Micromonospora* strains, core genes still harbored highly novel biosynthetic resources.

Overall, these results reflected strong stability and significant biosynthetic features of the genus *Micromonospora* in the production of Terpene and PKSother natural products. Even within the core genes, *Micromonospora* BGs remain highly novel, indicating the vast untapped resources of natural products within this genus. In addition, the extensive accessory genes represent an important reservoir of biosynthetic potential, reflecting the expanded biosynthetic capabilities of *Micromonospora* through lineage diversification. Furthermore, the unique BGs found in singleton genes suggested an ongoing evolutionary process where individual *Micromonospora* strains continuously acquire new biosynthetic capabilities.

### Metabolomic characteristics of *Micromonospora* isolates

To further explore the metabolic capabilities of the 15 isolates, fermentation experiments were conducted using two liquid cultures, named GYM and V-22. Each sample was subjected to UPLC-QTOF-MS/MS metabolomics analysis after proper incubation time. The data analysis revealed that a total of 33,999 features were obtained from the mass spectrometry raw data. Among them, 650 features were of high quality and contained MS/MS fragment ion pairs. The analysis resulted in the construction of feature based molecular network (FBMN) for the 15 *Micromonospora* isolates as shown in (**Fig. 4A**). Within the FBMN, there were 233 features assigned to Alkaloids (35.8% of the total), 164 features were assigned to Amino acids and Peptides (25.2% of the total), 117 features were assigned to Fatty acids (18% of the total), 25 features were assigned to Carbohydrates (3.8% of the total), 11 features were assigned to Terpenoids (1.7% of the total), 10 features were assigned to Polyketides (1.5% of the total), 6 features were assigned to Shikimates and Phenylpropanoids (0.9% of the total), and 84 features were categorized as Unknown (12.9% of the total).

**Fig. 4.**
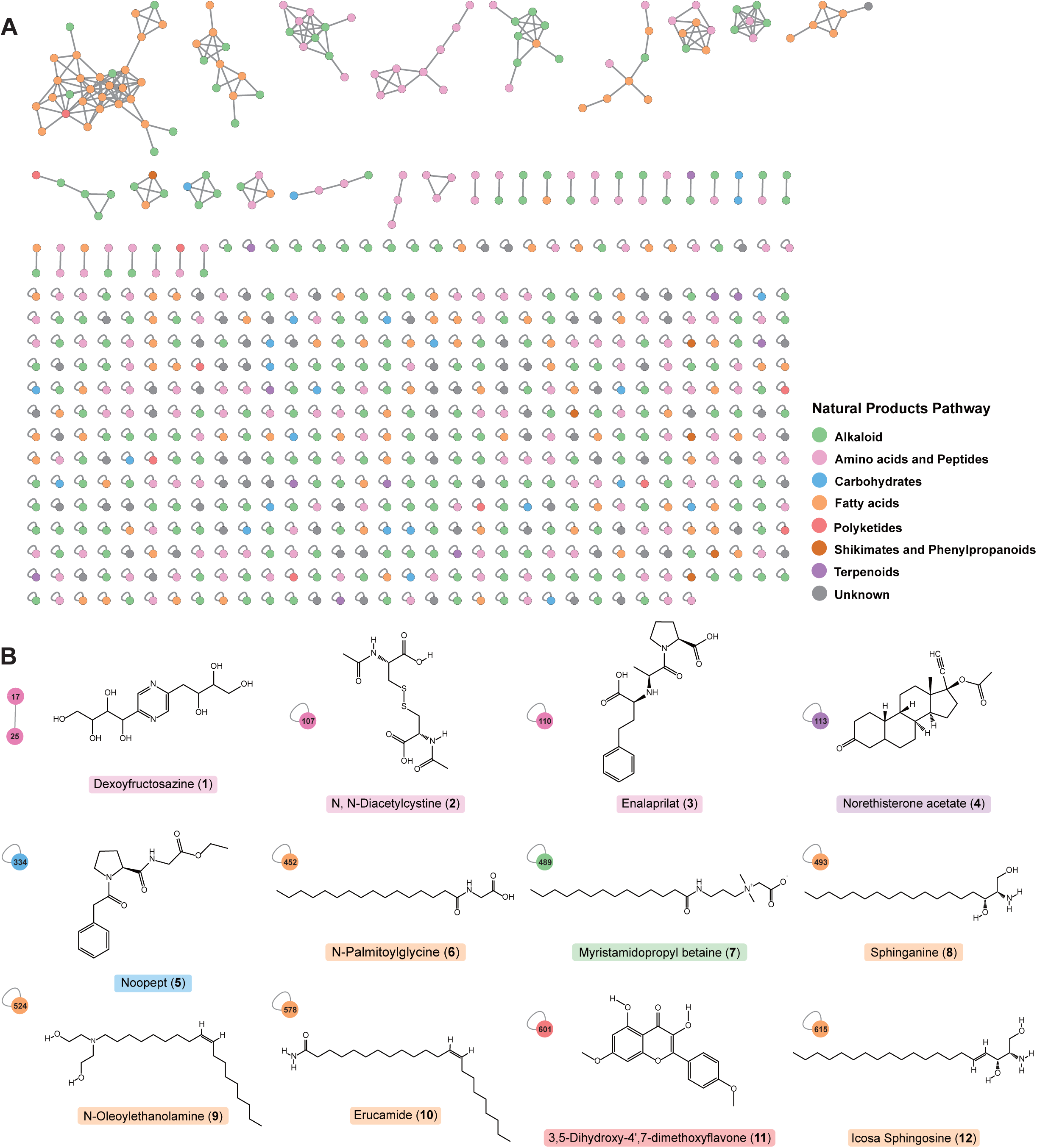
Metabolomic characteristics of *Micromonospora* isolates. **A** The feature-based molecular network (FBMN) constructed based on the UPLS-QTOF-MS/MS analysis of fermentation products from *Micromonospora* isolates. Each node in the network represents a mass spectral feature, and different colors indicate the identified natural product pathway categories. **B** The features were computed by CSI:fingerID using MS/MS mass spectrometry data to generate fragment trees, and then utilized machine learning to predict molecular structural fingerprints. These fingerprints were subsequently compared against the PubChem molecular structure database, resulting in matches with 12 known functional molecular structures.

Among the top matches of predicted molecular structures, 12 are known functional molecules, including Dexoyfructosazine, N, N-Diacetylcystine, Enalaprilat, Norethisterone acetate, Noopept, N-Palmitoylglycine, Myristamidopropyl betaine, Sphinganine, N-Oleoylethanolamine, Erucamide, 3,5-Dihydroxy-4’,7-dimethoxyflavone, and Icosa Sphingosine (**Fig. 4B**, **Table S6**). Moreover, as discussed in the early sections regarding the biosynthetic characteristic of genus *Micromonospora* is the biosynthesis of terpenoid and polyketide natural products, 20 features annotated as terpenoids and polyketides pathways in the metabolome study were observed that are of particular interest. Furthermore, among the top matches of predicted molecular structures, two known molecules were identified, including, Norethisterone acetate (compound **4**) and 3,5-Dihydroxy-4’,7-dimethoxyflavone (compound **11**). It is worth noting that compared to the biosynthetic potential at the genetic level, metabolomic data based on mass spectrometry provides a more direct reference value for natural product exploration, as these features are theoretically obtained through the isolation of fermentation products. These results revealed that the novel strains belong to *Micromonospora* genus contains genes to synthesis natural products.

### Relationship between phylogeny, biosynthetic genes, and metabolites in *Micromonospora*

As demonstrated in the previous results, the number of different classes of BGCs in *Micromonospora* varied significantly across phylogenetic clades (**Fig. 1C**). Furthermore, random forest models constructed based on the distribution of BGCs within GCFs also highlighted the contribution of phylogeny to the diversity of biosynthetic potential (**Fig. 2C**). To further investigate this, the study delved deeper into the relationships between phylogeny, biosynthetic genes, and metabolic profiles in *Micromonospora*. The analyses and results were presented as follows.

The hierarchical clustering trees via Jaccard and Bray-Curtis dissimilarity indices were constructed for GCFs and the metabolomic data to provide valuable insights into the evolutionary and functional relationships between the *Micromonospora* strains. Subsequently, the topological structures of these two clustering trees were compared with phylogenetic tree based on the genomes of *Micromonospora* isolates as shown in **Fig. 5A**. The results revealed that biosynthetic gene tree exhibited the highest correlation with the genomic phylogenetic tree (Pearson correlation = 0.709, Baker’s Gamma Index = 0.394, and *p* = 0.001 vs null model), thus confirming the previously mentioned correlation between phylogenetic lineage and differentiation of biosynthetic resources. On the other hand, a non-significant correlation was observed between the metabolomic tree and the genomic phylogenetic tree (Pearson correlation = 0.061, Baker’s Gamma Index = 0.137, and *p* = 0.145 vs null model). Similarly, there was also a non-significant correlation between the biosynthetic gene tree and the metabolomic tree (Pearson correlation = 0.066, Baker’s Gamma Index = 0.052, and *p* = 0.207 vs null model) as shown in **Fig. S6**.

**Fig. 5.**
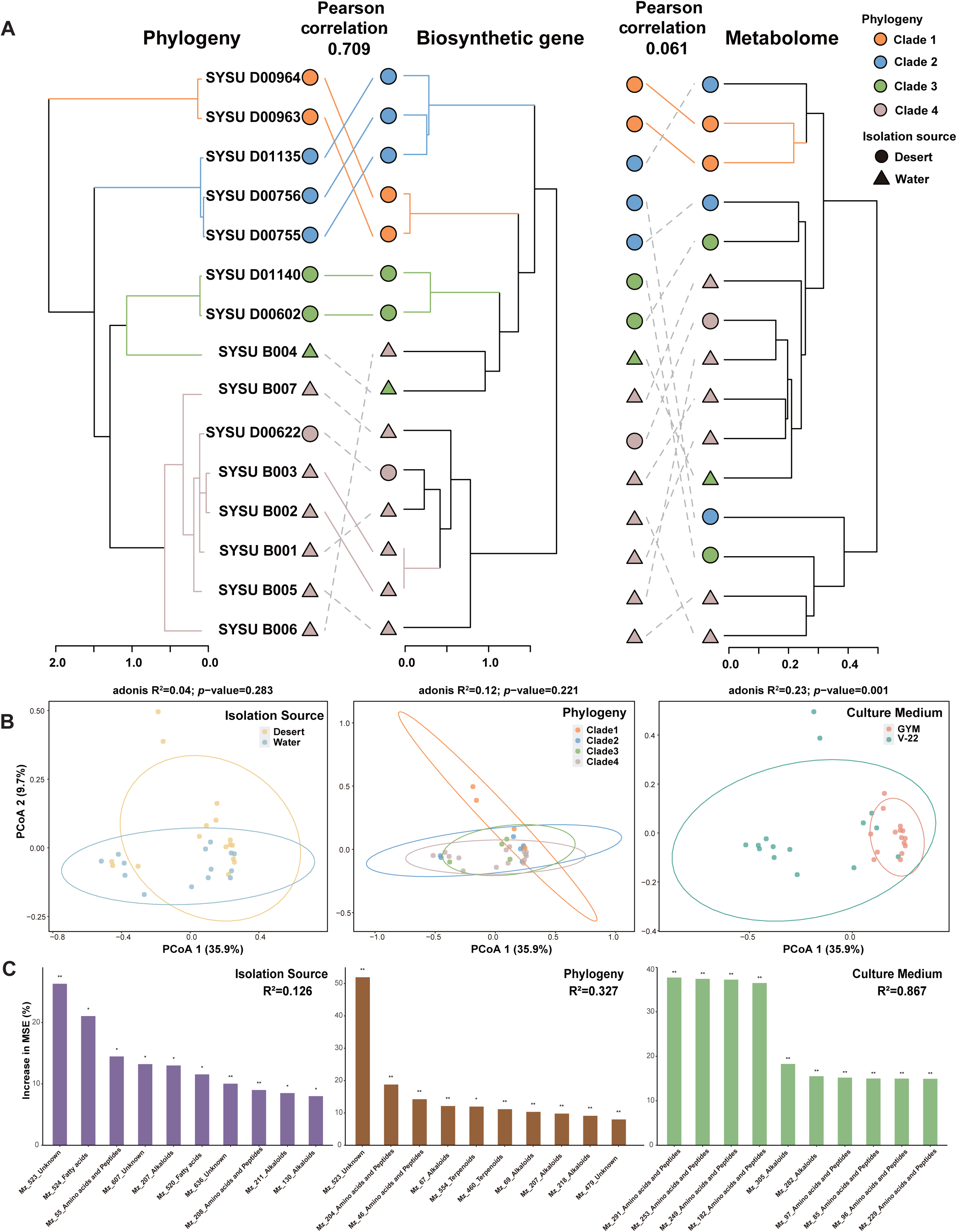
Comparative analysis of phylogeny, biosynthetic genes, and metabolites in *Micromonospora* isolates. **A** Hierarchical clustering trees of the biosynthetic genes and metabolomic data of *Micromonospora* isolates based on Jaccard and Bray-Curtis dissimilarities, respectively, compared with the phylogenetic tree constructed using genomes. Solid and colored lines connect matching subtrees in the two trees. **B** PCoA clustering analysis based on the metabolomic data of *Micromonospora* isolates, divided into three groups based on isolation sources, phylogenetic relationship, and culture media. **c** Random Forest (RF) models (5000 trees) constructed based on the metabolomic data, using isolation source, phylogeny, and culture medium as grouping criteria. The bar chart displays the top 10 features with the highest mean predictor importance (percentage of increase of mean square error) in each model. Percentage increases in the MSE (mean squared error) of variables were used to estimate the importance of these predictors, and higher MSE% values imply more important predictors. Significance levels are as follows: *p* < 0.5 = *, *p* < 0.01 = **. MSE, mean squared error.

Moreover, we calculated the distance between phylogenetic and biosynthetic genes and metabolomic data using differential trees and correlation analyses were conducted, as shown in **Fig. S7**. The results showed a significant positive correlation between phylogenetic distances and biosynthetic gene distances (R^2^=0.63). However, a non-significant correlation was observed between phylogenetic distances and metabolomic distances. Similarly, non-significant correlation was found between biosynthetic gene distances and metabolomic distance. These results highlight the complex relationship between phylogenetic, biosynthetic, and metabolomic diversity in the *Micromonospora* genus. The strong correlation between phylogenetic and biosynthetic gene distances suggests that the evolution of biosynthetic capabilities is closely tied to the overall genomic evolution of these strains. However, the lack of correlation between these factors and the metabolomic profiles indicates that additional factors, such as gene expression, regulation, and metabolic flux, may play important roles in shaping the observed secondary metabolite diversity.

Principal coordinate analysis (PCoA) was conducted based on the abundance of features with MS/MS fragment ion pairs as shown in **Fig. 5B**. The samples were divided into three groups based on differences in isolation source, phylogeny, and culture medium. PCoA analysis results demonstrate that the metabolomic data of *Micromonospora* isolates formed significant clusters only at the culture medium grouping level (ADONIS test). Clustering based on phylogeny showed a better fit and smaller probability (*p*-values, non-significant) compared to the clustering based on isolation source, indicating that PCoA clustering based on phylogeny non-significantly outperformed the clustering based on isolation source. Furthermore, random forest models were constructed using the same data and grouping criteria (**Fig. 5C**). The analysis revealed that the model based on culture medium grouping (R^2^=0.867) exhibited higher explanatory power and better fit compared to the models based on phylogeny (R^2^=0.327) and isolation source (R^2^=0.126). Moreover, the trend in the fit of the random forest models based on the metabolic data for the phylogeny and isolation source groups was consistent with the random forest model constructed based on the distribution of GCFs. In all cases, the model fit for the phylogeny grouping was higher than that for the isolation source grouping. These findings were interesting as they suggested that the relationship between phylogeny and isolation source governed the differentiation of biosynthetic resources in *Micromonospora* at the genomic level and was mirrored in the phenotype of metabolomic data. However, the phenotype of metabolomic data was most influenced by the newly introduced differences in culture medium as evidenced by the significant impact observed. These results provided insights into the evolutionary and environmental factors shaping natural product biosynthesis in the *Micromonospora* genus, with implications for bioprospecting and natural product discovery efforts targeting this group of *Actinobacteria*.

Therefore, the phylogenetic relationship in *Micromonospora* isolates was significantly correlated with the biosynthetic genes, but the metabolomic data was dissociated from both the genomic phylogeny and biosynthetic genes.

## Discussion

In this study, we isolated 15 *Micromonospora* strains from two distinct environments—marine sediment and desert soil—and combined them with reference genomes to assess the biosynthetic potential of a total of 107 *Micromonospora* genomes. A pan-genomic analysis was performed to characterize their biosynthetic features and distribution. Additionally, integrated genomic and metabolomic analyses of the 15 isolated strains were conducted to explore the correlations among phylogenetic relationships, biosynthetic capacities, and metabolic profiles. This study offered a comprehensive overview of the biosynthetic potential within the genus *Micromonospora*, providing novel insights into the exploration of microbial natural products through multi-omics strategies.

The advancements in genome-based bioprospecting technologies have revolutionized the way we approach natural product discovery and development. These approaches integrate molecular biological techniques to activate or heterologously express BGCs and employ bioinformatics tools to predict the structures of natural products encoded by BGCs(25). Subsequently, chemical synthesis enables researchers to obtain the target compounds(39). Thus, systematically assessing the biosynthetic capacities of previously characterized microbial genomes, as well as exploring uncharacterized microbial species, is critical to expanding genomic resources and enhancing natural product research. The analysis of 107 *Micromonospora* genomes revealed significant diversity and novelty in their BGC repertoires (**Fig. 2A**). Notably, several BGCs from the strains in this study clustered into GCFs along with reference BGCs known to encode bioactive compounds (**Fig. S6**). These findings underscored the potential of *Micromonospora* as a valuable resource for natural product discovery, particularly from strains inhabiting extreme environments.

The biosynthetic potential of microbes encoded in their genomes plays an important role in mediating resource competition within ecological systems(34). This potential, comprised of various BGs, is considered crucial for the consolidation and expansion of microbial niches(40, 41). The actinobacterial lineage, which includes the genus *Micromonospora*, represents a particularly ecologically adaptable group that thrives in diverse range of environments worldwide(42). Such adaptability is closely linked to the extensive metabolic repertoire possessed by these microorganisms (43). Despite this, the mechanisms underlying the formation and diversification of microbial biosynthetic capacities remain poorly understood. Insufficient knowledge of these mechanisms poses challenges to unlocking the natural resources embedded within wild-type microbial strains.

To address these gaps, this study conducted a detailed exploration of the biosynthetic potential of *Micromonospora* using pan-genomic analysis. The results demonstrated that the number of singleton genes in the novel *Micromonospora* isolates was stable across different pan-genome datasets (**Fig. 3B**), highlighting the stability of gene acquisition crucial to its biosynthetic capabilities. Within the pan-genome dataset (**Fig. 3A, 3C**), PKSother and Terpene BGs were highly conserved, occupying a significant portion of the core gene pool. These findings underscored the foundational biosynthetic characteristics shared by the *Micromonospora* lineage. Conversely, other BG classes, including NRPS, PKS-NRP_Hybrids, RiPPs, and Others classes of BGs were highly associated with accessory and singleton genes, suggesting niche-specific or strain-specific biosynthetic adaptations. Moreover, two types of BGs, NI-siderophore and NAGGN (N-Acetylglutaminylglutamine amide), were separated from the Others BG classes. Previous studies have reported that NAGGN plays an important role in the adaptation mechanism to osmotic stress(44), and its BGs are mainly distributed in core genes indicating its crucial role in the survival of *Micromonospora*. Likewise, NI-siderophore is also closely related to the strain’s survival, assisting in the accumulation of iron in the organisms(45). However, contrary to NAGGN, NI-siderophore BGs are mainly distributed in accessory genes, and are more diverse. Such diversity may stem from evolutionary pressures requiring structural diversification for iron acquisition under competitive microbial interactions(45, 46). The observed gene distribution further corroborates the adaptive differentiation of biosynthetic potential within *Micromonospora*.

Traditionally, natural product discovery has relied on the isolation of bioactive metabolites through activity-guided fractionation(47). Recently, the integration of omics data, including genomics and metabolomics, has significantly enhanced discovery efforts by decreasing rediscovery rates, guiding experimental work towards highly promising metabolites, and revealing their biosynthetic enzymatic pathways(48).

Among omics approaches, metabolomics is particularly valuable for natural product discovery, as metabolomic profiling detects molecular features that theoretically correspond to isolatable metabolites(49, 50). In this study, a metabolomic analysis of *Micromonospora* demonstrated its fundamental biosynthetic capacity for the production of terpenes and polyketides, with 20 features annotated as pathways related to these metabolites (**Fig. S7**). Through comparison with molecular structure databases using CSI:fingerID, two known molecules were identified: Norethisterone acetate and 3,5-Dihydroxy-4’,7-dimethoxyflavone. Furthermore, the metabolomic data also matched 12 known functional molecules (**Fig. 4B**), all possessing significant medicinal and industrial value (**Table S6**). Notably, these functional molecules were not previously reported as being isolated from *Micromonospora*. The biosynthetic potential and industrial application value of *Micromonospora* isolates, as evidenced by their capacity to produce structurally diverse and biologically active natural products, highlight the significance of this genus in the field of natural product discovery and its potential for future applications.

Previous studies have highlighted the heterogeneity between the phylogeny of microbial biosynthetic core genes and the phylogeny of their species, underscoring its importance in understanding microbial evolution and natural product discovery(51–53). This complexity suggests that researchers must take into account not only the phylogenetic relationships of species but also the metabolic diversity across strains when exploring natural product resources. In this study, we observed a significant correlation between the number of BGCs, their distribution across GCFs, and phylogenetic relationships (**Fig. 1C, 2C**). Further genomic analysis revealed that the biosynthetic potential repertoire of *Micromonospora* isolates is strongly associated with their phylogenetic positions (**Fig. 5A, S8**). Traditionally, phylogenetic relationships have been primarily based on conserved protein sequences in the genome(54). However, BGs, which play a dominant role in biosynthetic capabilities, are not among these conserved protein sequences. This finding is contrary to previous research conclusions that suggest limited or no correlation between the phylogenetic relationships and the biosynthetic capabilities of microorganisms(51–53). This insightful observation highlights the need for a more nuanced and multifaceted approach to understanding microbial evolution and natural product discovery. By considering alternative evolutionary mechanisms beyond just phylogenetic relationships, researchers can better navigate the complex landscape of microbial biosynthetic potential and unlock the full potential of *Micromonospora* and other microbial resources.

Furthermore, when comparing the metabolic data grouped by isolation source, phylogeny, and different culture mediums, the metabolic data showed significant clustering in the culture medium grouping, and a random forest model with high fit was obtained (**Fig. 5B, 5C**). This indicated that culture conditions play a dominant role in realizing the metabolic potential of a strain through laboratory fermentation. Importantly, it underscored a key bottleneck in the current development of natural product resources: the biosynthetic potential encoded in microbial genomes often remains underutilized in laboratory and industrial production environments(34). Building on this observation, we propose strategies such as the OSMAC approach could optimize culture conditions to enhance the strain’s metabolomic diversity. This may eventually expand the metabolic repertoire to better reflect the BGCs repertoire, establishing a more robust correlation with phylogeny.

However, this study used only two culture media, and further studies are required to explore this hypothesis in greater depth. Additionally, the focus on the genus *Micromonospora* restricts the generalizability of the findings. To improve their universality across microorganisms, future research should incorporate a broader range of taxa and more comprehensive datasets for robust analysis and validation.

In summary, this study expanded the taxonomic and genomic understanding of the *Micromonospora* clade, while elucidating the biosynthetic potential. Through pan-genomic analysis, the core biosynthetic capacities of this genus to produce terpene and polyketide secondary metabolites were unveiled. These findings also suggested that the biosynthetic capabilities of *Micromonospora* appear to have evolved during their adaptive evolution and lineage differentiation processes, offering preliminary insights into the formation and evolution of microbial biosynthetic potential. Furthermore, multi-omics analyses established correlations between phylogenetic relationships and biosynthetic pathways, emphasizing the significant role of *Micromonospora* in natural product discovery. This study highlighted the importance of seeking novel strains for targeted biosynthesis research aimed at deriving bioactive compounds with specific therapeutic or industrial applications. Integrating metabolomic data, diverse culture conditions, and an understanding of cell regulation will be crucial for realizing the full biosynthetic potential of these valuable microbial resources. This holistic approach can lead to the efficient discovery of novel natural products with diverse structures and bioactivities, ultimately contributing to the development of new therapeutic agents and other valuable applications.

## Materials and methods

### Sample collection and isolation cultivation

Strains SYSU B001, SYSU B002, SYSU B003, SYSU B004, SYSU B005, SYSU B006, and SYSU B007 were isolated from surface sediment samples (10 cm) collected at the Pearl River Estuary, Guangzhou, Guangdong, China (23° 6′ N 113° 18′E). The sediment samples were diluted 10^3^-fold with sterile distilled water, and 0.2 ml of the final suspension was spread on R_2_A agar(55) plates. The isolation plates were incubated at 28°C for 7 days. Pure colonies were obtained from the R_2_A agar at 28°C. The purified strains were preserved in a glycerol suspension (20%, v/v) and stored at -80°C for future use.

Strains SYSU D00602, SYSU D00622, SYSU D00755, SYSU D00756, SYSU D00963, SYSU D00964, SYSU D01135, and SYSU D01140 were isolated from sandy soil samples collected in the Gurbantunggut Desert, Xinjiang, China (45° 27′ N, 86° 40′ E; 391 m above sea level). To isolate the strains, 5 g of each soil sample was dissolved in 50 mL of sterile physiological saline (0.85% NaCl, w/v), and subjected to ultrasonic treatment at 45 kHz for 2 minutes. Then, the mixture was incubated at 28°C and 200 rpm/min for 1 hour using sterile 3 mm glass beads. Subsequently, supernatant of the mixture was diluted 10^3^-fold with sterile physiological saline, and 100 μl of the dilution was spread on minimal medium (MM) agar (56) plates supplemented with streptomycin (50 mg/L), naphthoic acid (50 mg/L), kanamycin (50 mg/L), and potassium dichromate (50 mg/L). The plates were incubated at 28°C for 1 to 4 weeks. Pure colonies obtained were transferred to GYM agar (Malt extract 10 g/L, Yeast extract 4 g/L, Glucose 4 g/L,

Trace salt 1 ml/L, CaCO_3_ 2g/L, Agar 18 g/L, pH 7.2 ± 0.2) plates for purification. The purified strains were preserved in a glycerol suspension (20%, v/v) and stored at -80°C for future use. The detailed experimental methods for morphology, physiology and chemotaxonomy of strains can be found in the supplementary method.

### DNA extraction, sequencing, and assembly

The extraction of isolated strains’ genomic DNA was followed by the method described by Li et al.(57) using the universal bacterial primers 27F and 1492R(58). The 16S rRNA gene amplification method was the same as described by Li et al.(59). The purified amplification fragments were cloned into the pEASY-T1 vector (TransGen Biotech) and transformed into Trans1-T1 chemically competent cells (TransGen Biotech). The selected clones were sequenced by GENEWIZ Co., Ltd. (Guangzhou, China). The obtained sequences were quality-checked and assembled using the SeqMan program (DNAStar v.7.1.0) and searched against the EzBioCloud server (https://www.ezbiocloud.net/) and the NCBI nucleotide database (https://www.ncbi.nlm.nih.gov/) using BLAST to identify related species sequences.

The draft genome of the strains was sequenced on the Illumina NovaSeq PE150 platform at Beijing Novogene Biotechnology Co., Ltd. The raw sequencing data of the genome were assessed for quality using FastQC software (v.0.11.8) (60) to evaluate the sequencing quality of the genome sequence. Based on the quality assessment results, the fastp software (v.0.23.0) (61) was used for quality control to obtain high-quality genome sequence data. The SPAdes software (v.3.14.1) (62) was employed for genome sequence assembly, and sequences with a length below 500 bp were removed from the genome assembly. The quality of the assembled genome was evaluated using the QUAST software (v.5.1.0rc1) (63), and the completeness and contamination of the genome were analyzed using the CheckM software (v.1.1.3) (64) . All obtained genomes in this study exhibited completeness > 90% and contamination < 5%, indicating that they were suitable for subsequent analysis.

### Genome dataset construction, phylogenetic analysis and BGCs annotation

Based on the NCBI GenBank database, a keyword search using “*Micromonospora*” and “Reference genomes” was performed, resulting in the retrieval of 106 genomes. The CheckM software (v.1.1.3) was utilized to analyze the completeness and contamination of the genomes. Genomes with completeness > 90% and contamination < 5% were selected. Utilizing the BacDive database, a manual search was conducted to retrieve the ecological source information of the required bacterial strains. The BacDive database (https://bacdive.dsmz.de/) was used to retrieve the isolation source information of the genomes, and some genomes were manually searched as necessary. As a result, a total of 92 genomes were obtained, possessing complete isolation source information and meeting the quality criteria. These genomes covered 92 valid published *Micromonospora* species. Combined with the 15 *Micromonospora* strains isolated in this study, these 92 genome sequences formed the *Micromonospora* genome dataset, comprising a total of 107 genomes for further analysis. The dataset of 92 valid published *Micromonospora* species obtained covered completely the most relevant species to the 15 *Micromonospora* strains isolated in this study. Consequently, the corresponding 16S rRNA sequences were downloaded from the LPSN (List of Prokaryotic Names with Standing in Nomenclature) database (https://lpsn.dsmz.de/) for systematic analysis.

The phylogenetic trees based on the genomes of this study were constructed based on the concatenation of 120 ubiquitous single-copy proteins using GTDB-Tk software (v.2.3.0) (54, 65). Average nucleotide identities (ANI) were calculated using fastANI (v.1.33), and digital DNA–DNA hybridization (dDDH) values were calculated with the Genome-to-Genome Distance Calculator 3.0 (https://ggdc.dsmz.de/). The maximum likelihood phylogenetic tree based on 16S rRNA was constructed using the Tamura-Nei model (66) in Mega X (67).

BGCs from 107 *Micromonospora* genomes were predicted and annotated using antiSMASH 7.0(68). Similarity information between the predicted BGCs and the encoded BGCs for known natural products was obtained according to the results of antiSMASH. The genome annotation files were further used with BiG-SCAPE software (v.1.1.5) (53) to determine the types of biosynthetic pathways and construct gene cluster families (GCFs). The GCFs were visualized using Cytoscape software (v.3.9.1) to generate clustering networks.

### Pan-genomic analysis

In this study, pan-genomic analysis was conducted following the anvi’o 7.1 pan-genome workflow(69). To simplify the header lines of the 107 genome FASTA files, the “anvi-script-reformat-fasta” program was used. The FASTA files were then converted into an anvi’o contigs database using “anvi-gen-contigs-database”. The HMM models were modified into hits within the contigs database using “anvi-run-hmms”. The program “anvi-run-ncbi-cogs” was employed to annotate genes in the contigs databases with functions from the NCBI’s Clusters of Orthologous Groups (COGs). We utilized “anvi-export-gene-calls” to export a table containing gene caller IDs, along with their corresponding start and stop nucleotide positions. The boundaries of gene clusters were determined by taking the starting nucleotide (5’ end) of the first biosynthetic gene and the ending nucleotide (3’ end) of the last biosynthetic gene, as presented in the antiSMASH results. These nucleotide positions served as the starting nucleotide to establish the boundaries of the gene clusters. By linking the gene caller IDs to the BGCs based on their start and stop positions, we classified genes within the boundaries of a given BGC as biosynthetic genes for natural product production. Subsequently, we assigned classifications and potential compound names to the biosynthetic genes, both of which were derived from the composition of the BGCs. The resulting table was imported back into the contigs database using the “anvi-import-functions” command for integration. The external genomic storage was created using the “anvi-gen-genomes-storage” to store DNA and amino acid sequences, along with functional annotations for each gene. With the genomic storage in place, we utilized “anvi-pan-genome” with the genomic storage database, employing the options “--use-ncbi-blast” and parameters “--mcl-inflation 10”. In the pan-genomic dataset, genes that occurred 107 times were identified as core genes, genes that occurred once were identified as singleton genes, and the remaining genes were identified as accessory genes. The “anvi-summarize” program was used to obtain detailed information about the pan-genomic dataset. Based on this, the distribution of biosynthetic genes within the pan-genomic dataset and the pan-genomic characteristics of the genus *Micromonospora* were determined.

### Fermentation and UPLS-QTOF-MS/MS analysis

30 μL of bacterial liquid was aspirated from the glycerol tube and streaked onto a plate to obtain single clones. Single clones were then transferred to GYM liquid medium (18×180 mm test tubes containing glass beads). The cultures were incubated at 220 rpm for 3-5 days, with daily observations made. Tubes with dense liquid cultures were selected, and a 1% inoculum was transferred to 50 mL of GYM seed medium for cultivation at 220 rpm until the logarithmic phase was reached.

For further cultivation, a 10% inoculum was transferred to 50 mL of GYM and V-22 fermentation flask medium, and the cultures were incubated at 220 rpm for 5-7 days. Subsequently, 40 mL of the fermentation broth was transferred to a conical flask, and an equal volume of acetone was added. The mixture was sonicated, allowed to stand for 1 hour, and then centrifuged to collect the supernatant. The remaining liquid was concentrated to 15 mL using a rotary evaporator followed by freeze-drying. The dried material was dissolved in 2 mL of methanol to obtain the crude extract of the fermentation broth.

The obtained crude extract from fermentation was diluted 10-fold with methanol to prepare the test samples. The samples were then centrifuged at 10,000 rpm in a centrifuge to obtain the samples ready for analysis. The analysis of untargeted metabolomics in the samples was performed using an ACQUITY UPLC I-Class PLUS System coupled with a Synapt G2-Si Quadrupole-Ion mobility-TOF Mass Spectrometer (Waters Corporation, Milford, MA, USA). Chromatographic separation was conducted on an ACQUITY UPLC HSS T3 Column (2.1×100 mm, 1.8 μm, 100 Å; Waters Corporation) maintained at 35 ℃. Each sample was injected with 1 μL, and the mobile phase consisted of (A) 0.1% formic acid in water and (B) 0.1% formic acid in acetonitrile. The chromatography was carried out at a flow rate of 0.3 mL/min using the following program: 0-1 min, 98% A; 1-6 min, gradient of 98-80% A; 6-12 min, gradient of 80-2% A;12-15.4 min, 2% A; 15.4-15.5 min, gradient of 2-98% A; 15.5-18 min, 98% A.

Mass spectrometry detection was performed using ESI+ mode. The capillary voltage was set at 2.5 kV, the source temperature was maintained at 120°C, and the sampling cone was set to 35. The desolvation temperature was set to 350°C, with a cone gas flow rate of 40 L/Hr, desolvation gas flow rate of 800 L/Hr, and nebulizer gas flow rate of 6.5 Bar. For the acquisition of MS/MS data, the DDA mode was employed, with collision energy set in a range of 10-25 eV for low energy and 25-45 eV for high energy collisions. The lockspray reference compound was leu-enkephalin.

### Untargeted metabolomics data annotation and analysis

The raw UPLC-MS/MS data files were converted to mzXML file format using ProteoWizard (70). The mzXML files were imported into MZmine3 (71) for data processing, following the workflow: Mass detection, For Ms_1_ detection, “Noise level” was set to 1.0E^3^, and for Ms_2_ detection, “Noise level” was set to 5.0E^1^; Chromatogram building, “ADAP Chromatogram Builder” (72) was selected, with the “Scan filter” parameter set to Ms_1_, “Minimum consecutive scans” set to 5, “Minimum intensity for consecutive scans” set to 5.0E^1^, and “m/z tolerance” set to 0.002 m/z; Deconvolution, “Baseline resolver” was selected, with the “MS/MS scan pairing” option checked, The “Min peak height” was set to 1.0E^3^, and the “Baseline level” was set to 1.0E^3^; Isotopes, “Isotopic peaks finder” was chosen, with “m/z tolerance” set to 0.0005 m/z, and “Maximum charge of isotope m/z” set to 3; Alignment, “Join aligner” was selected, with “m/z tolerance” set to 0.01 m/z, “Weight for m/z” set to 75, “Retention time tolerance” set to 0.2 min, and “Weight for RT” set to 25; Gap filling, “Peak finder” was chosen, with “Intensity tolerance” set to 5.0%., “m/z tolerance” set to 0.01 m/z, and “Retention time tolerance” set to 0.2 min; Export feature list, CSV files were selected for statistical analysis, when selecting “Export molecular networking files”, the “Filter rows option” was set to “ONLY WITH MS2”, and the resulting CSV file and MGF file were utilized for importing into GNPS (73) for feature-based molecular networking (FBMN) (74) construction and into SIRIUS software (75) for annotation.

## Acknowledgements

This work was financially supported by the National Natural Science Foundation of China (32270076), Guangzhou Basic and Applied Basic Research Foundation, China (2025A04J3495), Guangdong Province Modern Agricultural Industry Technology System Innovation Team Construction Project Focused on Agricultural Products (Agricultural Microorganisms) (2024CXTD04).

## Author information

### Contributions

JRH: formal analysis, visualization, writing—original draft. SL, WHL, LX, LD, GYS, QCW, JLL : sampling, morphological identification. MA, WJL: writing—review and editing. LD: supervision, conceptualization, writing—review and editing. All authors read and approved the final version of the manuscript.

## Ethics declarations

### Animal and human rights statement

This article does not contain any studies with human participants or animals performed by any of the authors.

